# Increasing temperature weakens the positive effect of genetic diversity on population growth

**DOI:** 10.1101/2021.05.13.444034

**Authors:** Alexandra L. Singleton, Samantha Votzke, Andrea Yammine, Jean P. Gibert

**Affiliations:** Duke University, Department of Biology, Durham, NC, USA

**Keywords:** Genetic variability, Intraspecific variation, Intraspecific variability, global warming, microbes

## Abstract

Genetic diversity and temperature increases associated with global climate change, are known to independently influence population growth and extinction risk. Whether increasing temperature may influence the effect of genetic diversity on population growth, however, is not known. We address this issue in the model protist system *Tetrahymena thermophila*. We test the hypothesis that at temperatures closer to the species thermal optimum (i.e., the temperature at which population growth is maximal), genetic diversity should have a weaker effect on population growth compared to temperatures away from the thermal optimum. To do so, we grew populations of *T. thermophila* with varying levels of genetic diversity at increasingly warmer temperatures and quantified their intrinsic population growth rate, *r*. We found that genetic diversity increases population growth at cooler temperatures, but that as temperature increases, this effect almost completely disappears. We also show that a combination of changes in the amount of expressed genetic diversity (G), plastic changes in population growth across temperatures (E), and strong GxE interactions, underlie this temperature effect. Our results uncover important but largely overlooked temperature effects that have implications for the management of small populations with depauperate genetic stocks in an increasingly warming world.

## INTRODUCTION

Rapid global climate change has a myriad of ecological consequences, from individuals to ecosystems[1–7]. Rising temperatures, in particular, influence metabolic rates[8, 9], which determine reproduction[10–12] and mortality[13, 14], thus setting demographics and population growth[10, 15]. As a consequence, species have thermal tolerances, and these thermal tolerances ultimately determine where on the globe –and under what environmental conditions– species may survive and reproduce[16, 17]. As temperatures increase globally, whether species will shift their geographic ranges[17], or instead go extinct[2], will be largely determined by these temperature tolerances[18].

Genetic diversity has long been known to reduce species extinction risk (e.g., [19]). For example, genetic diversity is negatively related to extinction risk in birds[20], low genetic diversity increases extinction risk in butterflies[21], while genetic rescue (i.e., introduction of new genetic variants) decreases extinction risk in mice[22] and pigmy possums[23]. Genetic diversity thus hedges against changing environmental conditions by increasing the chance that a population will have individuals with high rates of survival in novel environmental conditions. However, a combination of habitat fragmentation and shifting environmental conditions often lead to geographic range reductions (e.g., mountaintop species[2]), or crashes in population size[24]. Smaller population size or geographic range strengthens drift and reduces genetic diversity, leading to higher inbreeding depression and extinction risk[19]. While both genetic diversity and temperature are well-known to independently influence population growth(e.g., [9, 19]), whether increasing temperatures may alter the effect of genetic diversity on population growth and extinction risk is largely unknown.

Here, we address this issue in a model microbial system, the globally distributed protist *Tetrahymena thermophila*. We do so because the genetic makeup of these organisms can be easily manipulated[25], and because these organisms play an important role in the global carbon cycle that ultimately determines the pace of climate change (i.e., the microbial loop, [26–28]). In particular, we address 1) whether genetic diversity influences population growth in *T. thermophila*, 2) whether temperature influences that effect, and, 3) through what mechanisms. We hypothesize that lower genetic diversity depresses population growth (lower intrinsic growth rate, *r*) through inbreeding depression [19], while higher genetic diversity increases growth. We also hypothesize that the effect of genetic diversity should be weaker near the species-level thermal optimum owing to differences in how different genotypes grow across temperatures (Topt, Fig 1a, b): at Topt, most genotypes should reproduce relatively well, while away from Topt, some genotypes may perform increasingly poorly. Consequently, increasing genetic diversity may increase the chance of the population having genotypes that reproduce well at temperatures away from Topt, resulting in a higher intrinsic growth rate with increasing genetic diversity (Fig 1a, b; blue). Conversely, only a weak relationship between genetic diversity and *r*, if any at all, should be observed near or at Topt (Fig 1a, b; red).

**Fig 1:**
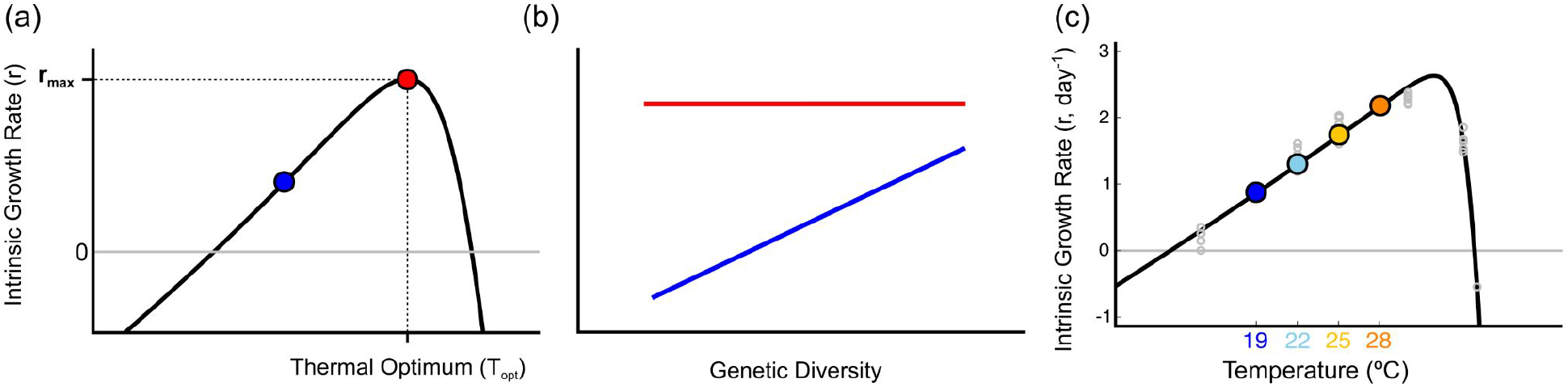
(a) Typical temperature performance curve for the population intrinsic growth rate, *r*, in solid black. Grey solid line represents r=0: above the line, the population grows, below, it decreases. We hypothesize that at temperatures away (blue dot) from the optimal temperature (Topt, red dot), increasing genetic diversity should lead to increasing intrinsic growth rate (b, blue solid line), while closer to the thermal optimum, increasing genetic diversity should not significantly increase *r* owing to similar growth rates across genotypes (b, red solid line). (c) *Tetrahymena pyriformis* temperature performance curve (black line, estimated from real data in grey). Colored dots indicate experimental temperatures (19°C, deep blue, 22°C, sky blue, 25°C, yellow, 28°C, orange).

## METHODS

### Experimental procedure

We sourced five clonal lines (B2086.2, A*III, CU438.1, A*V, and CU427.4) of the protist *Tetrahymena thermophila* from the Cornell University Tetrahymena Stock Center. The lines were reared in Carolina Biological protist medium® (Burlington, NC) in 200mL autoclaved borosilicate jars, and a 16-8 day/night cycle at 22°C within Percival growth chambers (Perry, IA).

To determine whether temperature alters the effects of genetic diversity on population growth, we manipulated the temperature and initial genetic diversity of microcosm populations. To manipulate genetic diversity, we started populations with a varying number of clonal lines (1, 2, 3, 4 or 5 lines). Monoclonal cultures were started with 50 individual protists. For all other treatments, the initial abundance of each clone depended on the total number of clones present, to control for possible effects of initial density: 2-clone populations started with 25 individuals/clone, 3-clone populations started with ~16 individuals/clone, 4-clone populations started with ~12 individuals/clone, and 5-clone populations started with 10 individuals/clone. Each monoclonal population, and each combination of four and five clones, was replicated 4 times. Each combination of two and three clones was replicated twice, for a total of 84 experimental populations per temperature. All experimental microcosms were reared in 3mL of growth media in 35 mm petri dishes.

We crossed the five genetic diversity treatments with four temperature treatments (19, 22, 25, 28 °C) along the rising portion of the temperature performance curve (TPC) of the species, for a total 336 experimental microcosms. The species TPC was quantified on a well-mixed population (Fig 1c). Experimental microcosms were grown in Percival growth chambers with all other environmental variables mimicking rearing conditions.

After a 24-hr incubation period, we estimated final population size through whole dish counts under a stereomicroscope (Leica, M205 C). Assuming exponential growth, the intrinsic growth rate (*r*) of each microcosm population was calculated as [log(Nf)-log(Ni)]/time, with time=1 day, *N_f_* being the final abundance, and *N_i_* the initial density (=50 ind for all experimental microcosms).

### Statistical Analyses

To test our hypotheses, we used a linear model with the log_10_ of *r* as the response variable and the log_10_ of the number of clones, temperature, and their interaction, as explanatory variables. To understand the mechanisms behind possible effects of temperature on the relationship between genetic diversity and *r,* we assessed whether changes in total additive genetic variation in *r* (G), environmental variation in *r* (E), or GxE interactions, could explain observed changes in *r* with genetic diversity and temperature. To do so, we used Analysis of Covariance (ANCOVA) on data from the monoclonal populations, with *r* as the response variable and temperature, clonal line, and their interaction, as explanatory variables.

## RESULTS

Population intrinsic growth rate (*r*) increased with temperature (estimate = 0.03±0.002, t = 20.70, p <0.001, Fig 2a) and increased with genetic diversity (estimate = 0.61±0.09, t = 6.94, p <0.001, Fig 2a). The positive effect of genetic diversity on *r* decreased with temperature (estimate = - 0.02±0.004, t = −5.60, p <0.001, Fig 2a). ANCOVA results show that this temperature influence on the effects of genetic variation on *r* is likely due to a combination of changes in the amount of expressed genetic variation in the populations, G, environmental changes in r, E, and strong GxE interactions (Table 1, Fig 2b).

**Table 1:** ANCOVA results assessing G, E, and GxE effects in *r*. Boldface indicates significant effects.

|  | Degree of Freedom | F-statistic | p-value |
| --- | --- | --- | --- |
| **G (differences across genotypes)** |  |  |  |
| At 19C° | 3 | 6.56 | **0.003** |
| At 22C° | 3 | 2.88 | 0.062 |
| At 25C° | 3 | 34.6 | **<10-6** |
| At 28C° | 3 | 2.88 | 0.059 |
| **E (differences across temperatures)** |  |  |  |
| Clone B2086.2 | 4 | 118 | **<10-8** |
| Clone CU427.4 | 4 | 26.8 | **<10-5** |
| Clone A*III | 4 | 48.3 | **<10-7** |
| Clone CU438.1 | 4 | 42.9 | **<10-6** |
| Clone A*V | 4 | 32.9 | **<10-6** |
| **GxE (differences across genotypes across temperatures)** | 12 | 4.70 | **<10-5** |

**Fig 2:**
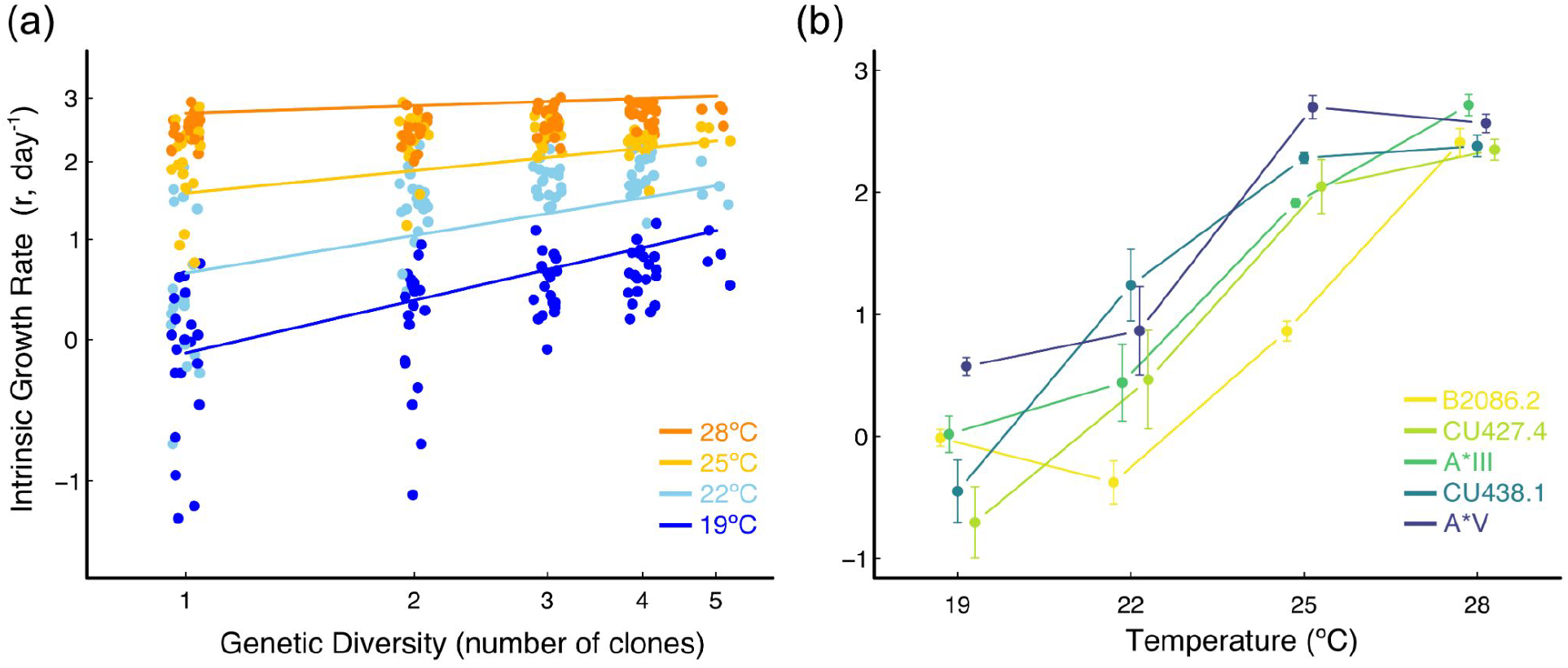
(a) Plot of the intrinsic growth rate, *r*, against the number of clones for all 336 experimental microcosms (dots), across all four experimental temperatures. Solid lines represent linear model predictions (color code as in Fig 1c). (b) Intrinsic growth rate against temperature for all monoclonal cultures. Bars represent standard errors and lines connecting dots indicate changes in r across temperatures (B2086.2 is Yellow, CU427.4 is Yellow green, A*III is Green, CU438.1 is Blue green, A*V is Blue).

## DISCUSSION

Rapidly changing environmental conditions and genetic diversity are both well-known to independently influence population growth and extinction risk (e.g., [2, 3, 23, 29]). Whether rapid climate change may alter how genetic diversity influences population growth, however, is not known. Our results indicate that as temperature increases toward a species’ thermal optimum, genetic diversity has a weaker effect on the intrinsic population growth rate (Fig 2a). These results imply that the effect of genetic diversity on population growth is contingent on environmental conditions.

Our results also indicate that changes in expressed additive genetic variation in *r* (G) are responsible for lower levels of variation observed at higher temperatures, compared to those observed at lower temperatures (Fig 2b, Table 1), while plasticity (E) is responsible for the observed increase in population growth with temperature (Fig 2b). Moreover, strong GxE effects, where different genotypes grow differentially at different temperatures, underlie the weakening of the positive effect of genetic variation on population growth rate (Fig 2a): clonal lines grow at similar rates at warmer temperatures, but do so at distinctly higher or lower rates in colder temperatures (Fig 2b). The presence of strong GxE effectively shifts which genotypes grow better at different temperatures, indicating the possibility for temperature-mediated clonal sorting in these microbial populations. Rapid evolutionary change has been suggested as a possible mechanism through which organisms may fend off the negative impacts of climate change[30–33]. In line with these studies, our results suggest that rapid evolutionary change (in this case, through clonal sorting), may occur in species where different genotypes display different thermal responses (GxE).

Together, our results indicate possible ways in which increasing temperatures associated with climate change and depauperate genetic stocks resulting from habitat fragmentation may jointly affect population growth and extinction risk. We show that temperature and genetic diversity interactively influence population growth: populations with higher genetic diversity have a weaker response to temperature compared to genetically depauperate populations (Fig 2a). As a consequence, while genetic diversity hedges against increasing temperatures, inbred –or small– populations, may respond more strongly. These results have important implications for the management of threatened and other species of interest in a changing world.

## ACKNOWLEDGMENTS

We thank Zeyi Han for providing help and expertise with *T. thermophila* clonal lines and Kathleen Donohue for early discussions. SV, AY and JPG were supported by a U.S. Department of Energy, Office of Science, Office of Biological and Environmental Research, Genomic Science Program Grant under Award Number DE-SC0020362 to JPG.

